# Corticosteroids and cellulose purification improve respectively the in vivo translation and vaccination efficacy of self-amplifying mRNAs

**DOI:** 10.1101/2020.08.26.268706

**Authors:** Zifu Zhong, Séan Mc Cafferty, Lisa Opsomer, Haixiu Wang, Hanne Huysmans, Joyca De Temmerman, Stefan Lienenklaus, João Paulo Portela Catani, Francis Combes, Niek N. Sanders

## Abstract

Synthetic mRNAs are an appealing therapeutic platform with multiple biomedical applications ranging from protein replacement therapy to vaccination. In comparison to conventional mRNA, synthetic self-amplifying mRNAs (sa-mRNAs) are gaining increased interest due to their higher and longer-lasting expression. However, sa-mRNAs also elicit an innate immune response, which may complicate the clinical translation of this platform. Approaches to reduce the innate immunity of sa-mRNAs have not been studied in detail. In this work we investigated the effect of several innate immune inhibitors and a novel cellulose-based mRNA purification approach on the type I interferon (IFN) response, translation and vaccination efficacy of our formerly developed sa-mRNA vaccine against Zika virus. Among the investigated inhibitors, we found that topical application of clobetasol at the sa-mRNA injection site was the most efficient in suppressing the type I IFN response and increasing the translation of sa-mRNA. However, clobetasol prevented the formation of antibodies against sa-mRNA encoded antigens and should therefore be avoided in a vaccination context. Residual dsRNA by-products of the *in vitro* transcription reaction are known inducers of immediate type I IFN responses. We additionally demonstrate drastic reduction of these dsRNA by-products upon cellulose-based purification, consequently reducing the innate immune response and improving sa-mRNA vaccination efficacy.

## Introduction

Synthetic mRNAs have become an appealing therapeutic platform with multiple biomedical applications ranging from protein replacement therapy to vaccination [1, 2]. Compared to plasmid DNA and viral vectors synthetic mRNAs hold some important advantages. First, they do not have to cross the nuclear barrier to exert their function, making them effective in both dividing and non-dividing cells [1, 3]. Furthermore, synthetic mRNAs allow cell-free production and exert a transient and more predictable expression [1, 2]. In recent years, synthetic self-amplifying mRNAs (sa-mRNAs) have also been gaining interest because of their higher and longer-lasting expression compared to non-amplifying mRNAs [4, 5]. Self-amplifying RNAs encode an RNA-dependent RNA polymerase (replicase) that gives them the capacity to trigger a temporal amplification of their backbone. Additionally, this replicase also generates many copies of smaller “subgenomic RNA(s)” that encode the protein(s) of interest. By using synthetic sa-RNAs it is hence possible to reduce the dose and the need for repeated injections, while still benefiting from the desirable features of synthetic mRNAs.

However, innate immunity triggered by sa-mRNA may complicate the clinical translation of this platform. The current *in vitro* production process of synthetic (sa)-mRNAs generates by-products such as short abortive transcripts and double stranded (ds) RNA species, which are recognized as non-self by toll-like receptors (TLRs), cytoplasmic RIG-I like receptors (RLRs) and other cellular pattern recognition receptors (PRRs) [1, 6, 7]. This triggers the production of proinflammatory cytokines and type I interferons (IFN), which are undesirable when synthetic (sa)-mRNAs are considered for protein (replacement) therapy [7]. In contrast, the cytokines induced by this self-defence mechanism may serve as adjuvants and hence facilitate the effects of synthetic (sa)-mRNA vaccines [8]. However, this view needs to be nuanced as studies demonstrated that, depending on the administration approach, type I IFN responses can also decrease the efficacy of mRNA vaccines by negatively affecting immune responses [9] and by inducing enzymes that inhibit mRNA translation [10, 11]. An important breakthrough was achieved when Kariko et al. demonstrated that the innate immunity of synthetic mRNAs can be drastically reduced by incorporation of modified nucleotides and by reversed-phase high-performance liquid chromatographic (HPLC) purification [12-14]. However, this was only demonstrated for non-amplifying mRNAs. We recently demonstrated that the innate immune response triggered by the self-amplifying mRNA platform [4, 5, 11] reduced the cellular and humoral responses of an sa-mRNA vaccine against Zika virus [15]. Therefore, strategies that can reduce the innate immunity of sa-mRNAs may improve the efficacy of sa-mRNA vaccines and the acceptance of sa-mRNA therapeutics in general. Tempering the innate immunity of sa-mRNA by inclusion of modified nucleosides is expected to have a negative impact on the replication of the sa-RNA [4, 16]. Moreover, the large size (> 10 kb) of sa-RNAs makes them more prone to shearforces, which complicates their purification by reversed-phase HPLC. Therefore, alternative strategies to decrease the innate immunity of sa-RNAs are needed.

In this work we investigated the capacity of several innate immune inhibitors to temper the innate immune response and improve the expression of a self-amplifying mRNA vaccine against Zika virus (ZIKVac-sa-mRNA). The tested inhibitors (Fig. 1) were either mixed with the sa-mRNA vaccine or locally administered at the intradermal injection site. Local application of clobetasol caused the strongest reduction of the innate immunity and drastically improved the *in vivo* translation of the sa-mRNA. In an alternative approach to mitigate innate immune responses triggered by dsRNA by-products, we also purified the ZIKVac-sa-mRNA by a new cellulose-based procedure. This new purification process drastically reduced the innate immunity, improved the expression and vaccination efficacy of our ZIKVac-sa-mRNA vaccine.

**Fig. 1.**
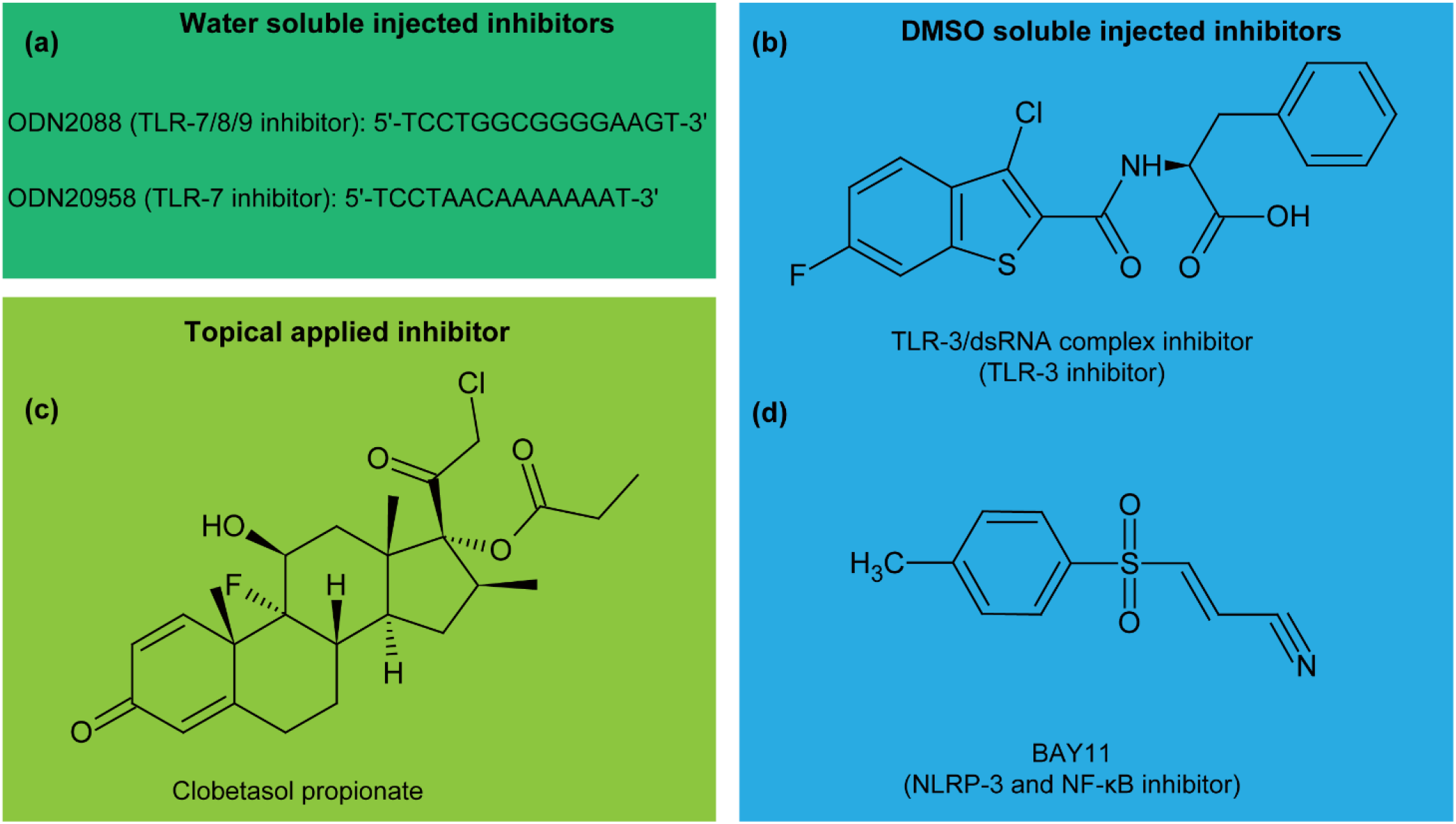
Structure of the innate inhibitors used in this study. Oligonucleotides ODN2088 and ODN20958 respectively inhibit TLR7/8/9 and TLR7 (a). Chemical structure of the TLR3/dsRNA complex inhibitor (termed “TLR3 inhibitor” in this paper) (b). Clobetasol propionate is a topically applied corticosteroid with broad immunosuppressive properties including NF-κB inhibition (c). BAY11 inhibits the intracellular NRLP3 receptor and NF-κB activation (d).

## Results

### Tempering the innate immunogenicity of self-amplifying mRNA with innate immune inhibitors

It is known that the *in vivo* safety and efficacy of sa-mRNA therapeutics is compromised by their strong activation of type I interferons (IFNs). To address this issue, we evaluated the capacity of a series of innate immune inhibitors (Fig. 1) to suppress the *in vivo* type I IFN response elicited by an sa-mRNA vaccine against Zika virus (ZIKVac-sa-mRNA). To this end, we co-administered innate immune inhibitors with the ZIKVac-sa-mRNA in the skin of IFN-β luciferase reporter mice (IFN-β^+/Δβ-luc^ mice) [17]. In these transgenic mice the luciferase expression is under control of the promotor of IFN-β, a key type I IFN. In the absence of co-administered innate immune inhibitors the IFN-β expression rapidly increased and peaked within 0-5 h after intradermal electroporation of the ZIKVac-sa-mRNA vaccine. After this peak the IFN-β expression sharply dropped and the background IFN-β expression was reached after about one week (Fig. 2a). Furthermore, we confirmed that the elicited IFN-β response is mainly occurring from the ZIKVac-sa-mRNA vaccine, as electroporation of solely PBS induced only a moderate type I IFN response (Fig. 2a, black curve). Co-injection of the water-soluble oligonucleotide-based TLR inhibitors ODN2088 or ODN20958 (Fig. 1a) with the ZIKVac-sa-mRNA vaccine significantly reduced the immediate type I IFN response (Fig. 2b-c). However, this innate immune tempering effect was lost after one day. The inhibitory effect of ODN2088, which blocks TLR7/8/9, was slightly higher than that of ODN20958, which only blocks TLR7 (Fig. 2b-c). A lower, but still significant, reduction of the early IFN-β response was also achieved when the ZIKVac-sa-mRNA was co-injected with lower doses (< 20 µg) of these TLR inhibitors (Fig. S1 a-b).

**Fig. 2.**
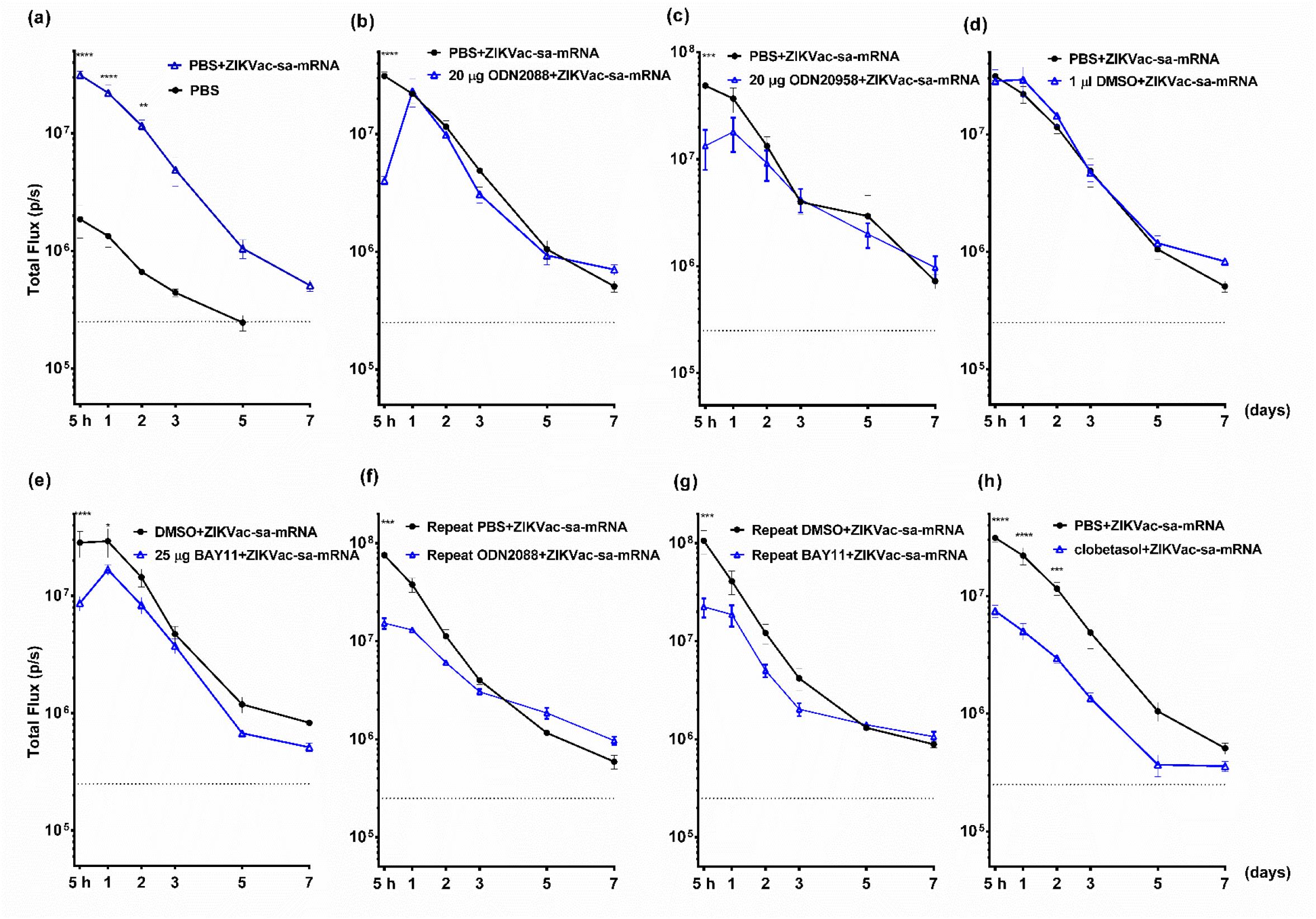
Effect of innate immune inhibitors on the kinetics of the type I IFN response after intradermal electroporation of a sa-mRNA vaccine against Zika virus in IFN-β^+/Δβ-luc^ reporter mice. Y-axis values represent the type I IFN response after a single intradermal electroporation of the ZIKVac-sa-mRNA vaccine (1 µg in 50 µl) or PBS control (a) and the capacity of inhibitors ODN2088 (b, f), ODN20958 (c), BAY11 (e, g) and clobetasol (h) to temper the innate immunogenicity over 1 week. Inhibitors ODN2088, ODN20958, BAY11 were mixed with the ZIKVac-sa-mRNA vaccine before administration. Since DMSO is needed to dissolve BAY11, the influence of DMSO was also studied (d). Panels (f) and (g) depict the effect of repeated ODN2088 or BAY11 administration where the injection site was first intradermally injected with the inhibitors 5 hours prior to ZIKVac-sa-mRNA administration, followed by a second inhibitor treatment 7 hours later. Local injection of the inhibitors continued twice daily until day 5. In contrast to the other inhibitors, clobetasol propionate was topically applied (25 µg in 1 cm^2^) one day prior to ZIKVac-sa-mRNA administration and repeated twice daily until day 3. Each symbol represents the mean of four individual mice and the error bars represent SEM.

Next, we evaluated the TLR-3/dsRNA complex inhibitor and BAY11 (Fig. 1b-d). The latter inhibits the intracellular NOD-like receptor pyrin 3 (NLRP3) and nuclear factor kappa-light-chain-enhancer of activated B cells (NF-κB). DMSO was used to dissolve these inhibitors as both are water insoluble. We first confirmed that addition of small amount of DMSO (1 µl) to our ZIKV-sa-mRNA vaccine (50 µl) did not change its IFN-β induction capacity (Fig. 2d). Co-injection of ZIKV-sa-mRNA with 25 µg BAY11 significantly suppressed the IFN-β response during the first 24 h (Fig. 2e), while 12.5 µg of BAY11 was not effective (Fig. S1 c). In contrast, neither of the tested TLR-3 inhibitor doses tempered the intrinsic innate immunogenicity of the ZIKVac-sa-mRNA vaccine (Fig. S1 d-e).

In a subsequent experiment we studied whether pretreatment and posttreatment of the injection spot with BAY11 or ODN2088 could increase and prolong their capacity to temper the innate immunogenicity of our sa-mRNA vaccine. Surprisingly, pretreatment of the injection site with BAY11 or ODN2088 and twice daily intradermal administration of these inhibitors after the injection of the ZIKVac-sa-mRNA did slightly, but not drastically increase or prolong the suppression of the IFN-β response (Fig. 2f-g). In another attempt to quell the type I IFN response, we considered topical application of the corticosteroid clobetasol. To that end, the injection site was pretreated with a clobetasol ointment 12 h prior to administration of ZIKVac-sa-mRNA. Clobetasol treatment was subsequently repeated twice daily during three days starting at the day of ZIKVac-sa-mRNA injection. This schedule of local clobetasol treatment drastically reduced and shortened the elicited type I IFN response (Fig. 2h). A significant inhibitory effect was observed up to two days after ZIKVac-sa-mRNA injection and overall a 3-fold reduction of the IFN-β expression was observed (Fig. 2i).

### Co-administration of multiple innate immune inhibitors

We next evaluated whether co-administration of multiple innate immune inhibitors could further decrease the type I IFN response elicited by our ZIKVac-sa-mRNA vaccine. In more detail, clobetasol was co-administered with either ODN2088 or BAY11, or with both inhibitors. In these experiments clobetasol was applied locally at the injection site twice daily for three days starting from the day of ZIKVac-sa-mRNA injection. The ODN2088 or BAY11 inhibitors were given as a single co-injection with the ZIKV-sa-mRNA (Fig. 3a). The clobetasol pretreatment, which was skipped in this experiment, seems to be of great importance as the inhibition of the IFN-β response was much lower without pretreatment of clobetasol (Fig. 2h and Fig. S1 f versus Fig. 3b and Fig. 3f). A drastic reduction of the type I IFN response was observed when clobetasol was combined with BAY11 or/and ODN2088 (Fig. 3c-e and Fig. 3g-i). Especially, the combination of the three inhibitors (clobetasol+ODN2088+BAY11) was very successful in inhibiting the type I IFN response elicited by our sa-mRNA vaccine (Fig. 3e and 3i).

**Fig. 3.**
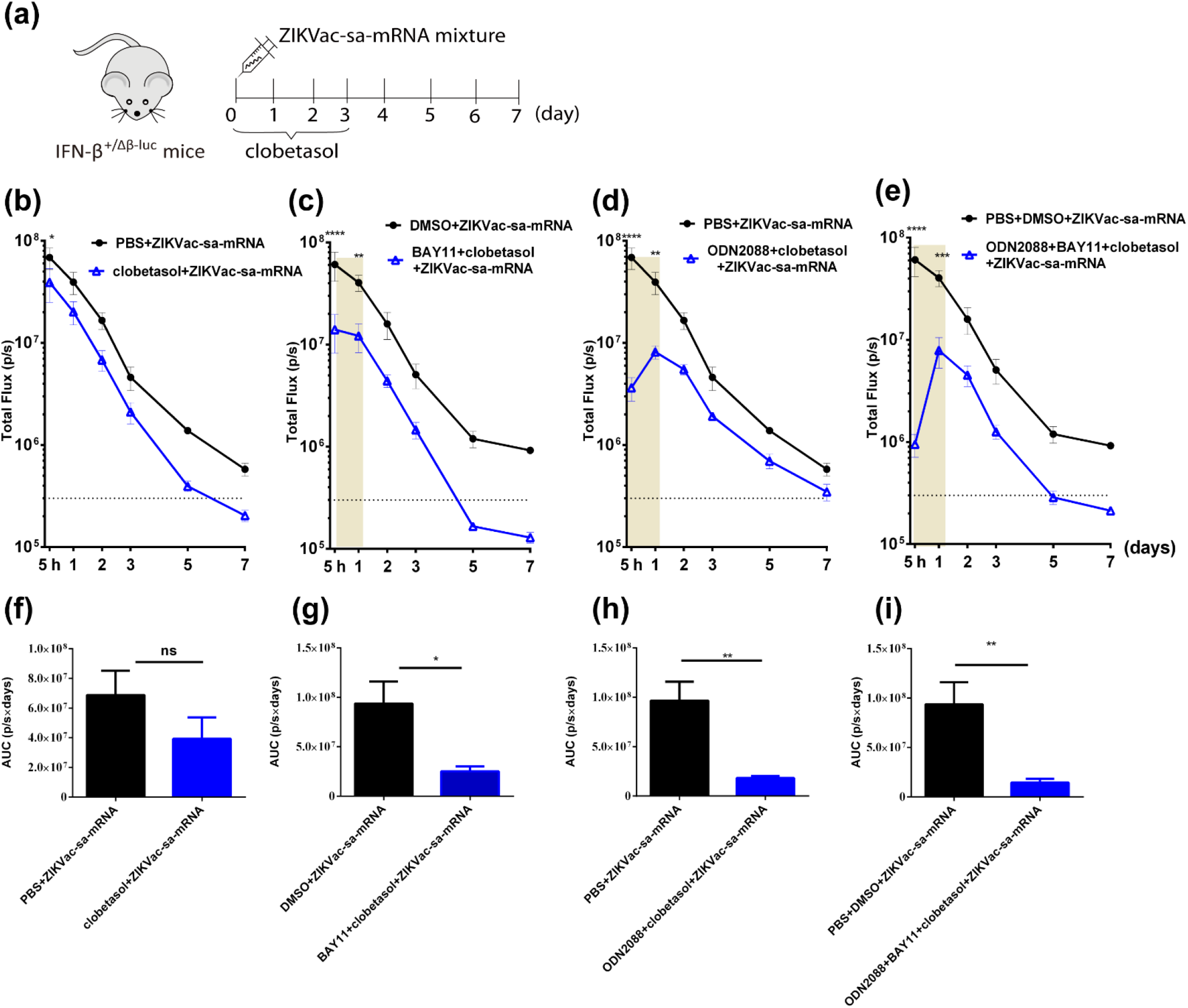
Effect of topical clobetasol in combination with other inhibitors on the type I IFN response kinetics after intradermal electroporation of ZIKVac-sa-mRNA in IFN-β^+/Δβ-luc^ reporter mice. Treatment schedule of the injection site (a). ZIKVac-sa-mRNA vaccine (1 µg in 50 µl) administration was directly followed by topical application of clobetasol propionate (25 µg in 1 cm^2^) twice per day and continued over 3 days (b). Additionally, clobetasol was combined with 25 µg BAY11 (c), 20 µg ODN2088 (d), or ODN2088 and BAY11 together (e). These inhibitors were co-injected once with the ZIKVac-sa-mRNA. The area under the curves (AUCs) of graphs b-e are presented in f-i, respectively. Each symbol or bar represents the mean of four individual mice and the error bars represent SEM.

### Influence of innate immune inhibitors on the translation of self-amplifying mRNA

It is generally accepted that a strong innate immune response after mRNA delivery has a negative impact on its translation efficacy [9-11, 15]. Therefore, we evaluated in BALB/c mice the effect of clobetasol, BAY11 and ODN2088 on the translation efficacy of sa-mRNA encoding luciferase (LUC-sa-mRNA). The LUC-sa-mRNA was again administered by intradermal electroporation. Pretreatment and posttreatment of the LUC-sa-mRNA injection site twice daily with a clobetasol ointment during 3 days prolonged the translation with one week and increased the initial translation within the first 6 days (Fig. 4a-b). This treatment regimen with clobetasol caused a 3.5–fold increase in overall translation of the LUC-sa-mRNA (Fig. 4b and Fig. S2 a). In contrast, co-injection of LUC-sa-mRNA with ODN2088 did not change the translation profile (Fig. 4c). Surprisingly, co-administration of BAY11 drastically reduced the translation of our LUC-sa-mRNA (Fig. 4d and Fig. S2 b). This was not due to the DMSO solvent, as LUC-sa-mRNA with equal amounts of PBS and DMSO was as effective as LUC-sa-mRNA with only PBS (Fig. S2 c).

**Fig. 4.**
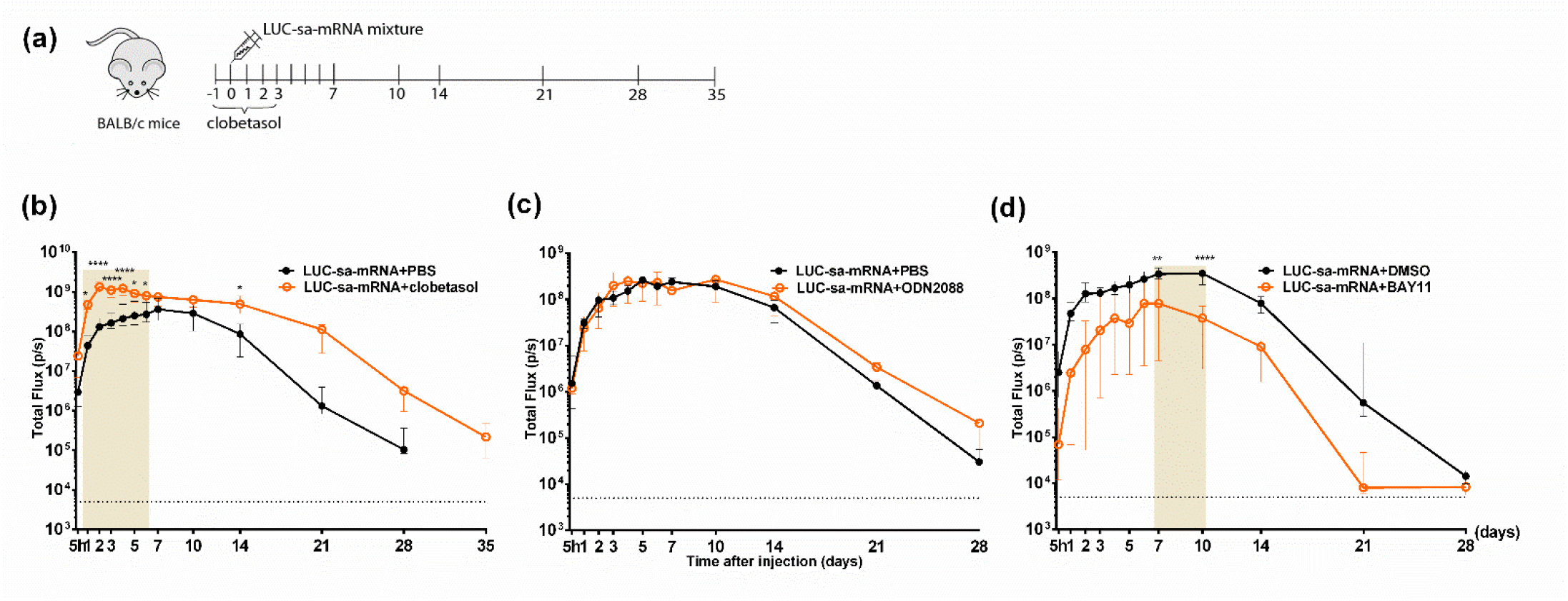
Influence of innate immune inhibitors on the translation of sa-mRNA encoding luciferase in BALB/c mice. Time schedule of the animal experiment (a). Wild type BALB/c mice were intradermally electroporated with 1 µg of sa-mRNA encoding luciferase (LUC-sa-mRNA) in the presence of clobetasol (b), ODN2088 (c) or BAY11 (d). Topical treatment of the injection site with clobetasol (25 µg in 1 cm^2^) was performed 1 day prior to LUC-sa-mRNA injection and subsequently twice daily for three days. BAY11 (25 µg) and ODN2088 (20 µg) were co-injected with the LUC-sa-mRNA (1 µg in 50 µl). The luciferase expression was determined by measuring the bioluminescent signal at the injection spot for 28 or 35 days. Each symbol represents the median of four individual mice and the error bars indicate interquartile range.

### Innate immunogenicity and translation efficacy of self-amplifying mRNA purified by cellulose chromatography

It is well-known that double-stranded (ds) RNA contaminants in synthetic (IVT) mRNAs play an important role in the activation of type I IFNs and translational inhibition [1, 18]. The classic purification strategies, like the silica-based columns which we routinely use, do not efficiently remove dsRNAs. However, recently it has been shown that these dsRNAs can be removed from short non-amplifying synthetic mRNAs by a cellulose-based purification method [18]. The applicability of this method to synthetic self-amplifying mRNAs, which are 3 to 4 times longer than non-amplifying mRNAs, is unknown. Therefore, in this section the capacity of this method to remove dsRNAs, to decrease the innate immunity and to improve the translation and vaccination efficacy of synthetic sa-mRNAs was investigated. We first demonstrated that synthetic sa-mRNAs purified with silica columns contained substantial amounts of dsRNA contaminants that can be efficiently removed by the novel cellulose-based purification method (Fig. 5a). Subsequently, the effect of cellulose-based purification on the innate immunity of the ZIKVac-sa-mRNA vaccine and the expression of LUC-sa-mRNA was studied after intradermal electoporation in IFN-β reporter and BALB/c mice, respectively. In addition, the effect of pre- and posttreatment (twice daily for three days) of the injection site with clobetasol was also studied. As shown in Fig. 5b, we confirmed that the silica-purified ZIKVac-sa-mRNA vaccine elicits a strong type I IFN response that can be tempered by clobetasol (blue and black curves). Interestingly, a similar reduction of the type I IFN response could be obtained when the ZIKVac-sa-mRNA vaccine was purified by the cellulose-based method (green curve). Topical application of clobetasol could not further decrease the overall elicited type I IFN response of the cellulose-purified ZIKVac-sa-mRNA (Fig 5c), albeit that clobetasol strongly reduced the IFN-β expression (i.e. circa 10-fold) 5 h after administration of the cellulose-purified ZIKVac-sa-mRNA vaccine (Fig. 5b red cruve and Fig. 5c). Although cellulose purification of the sa-mRNA significantly reduced the elicited type I IFN response, it only slightly improved the translation efficacy of the sa-mRNA (Fig. 5d, black and green curves). Nevertheless, co-administration of clobetasol significantly increased the translation of the silica-purified as well as cellulose-purified LUC-sa-mRNA. The highest expression was observed in the mice that received the cellulose-purified LUC-sa-mRNA together with clobetasol (Fig. 5d-e). As also observed in Fig. 4a, clobetasol seems to prolong the translation of both the silica- and cellulose-purified LUC-sa-mRNA (Fig. 5d).

**Fig. 5.**
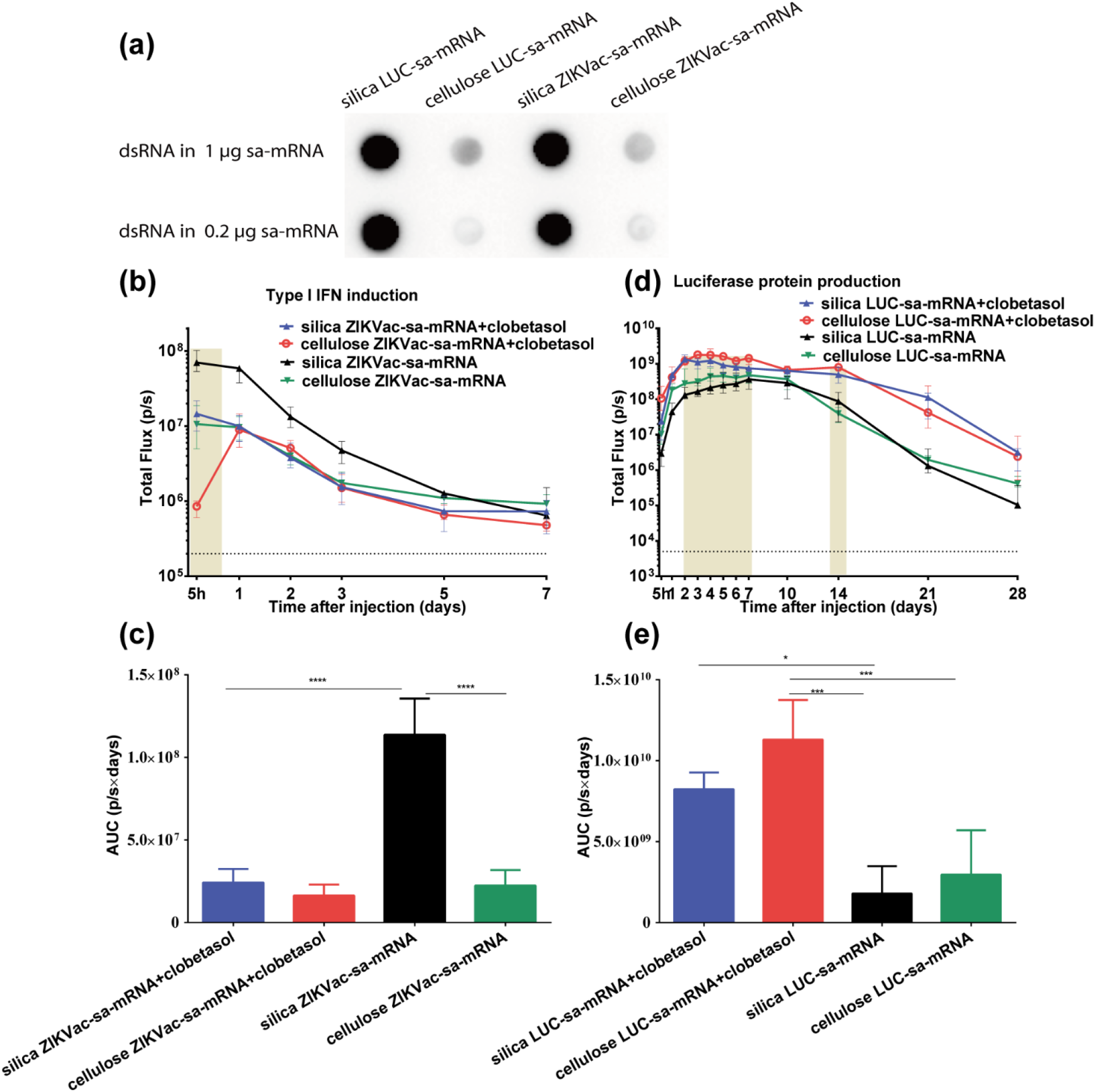
The effect of cellulose purification and clobetasol on the type I IFN response and translation of sa-mRNAs. dsRNA by-products in silica- and cellulose-purified sa-mRNAs (1000 or 200 ng µg per dot) as analyzed by dot blotting with J2 dsRNA-specific mAb (a). Type I IFN response kinetics after intradermal electroporation of silica- and cellulose-purified ZIKVac-sa-mRNA (1 µg) in IFN-β^+/Δβ-luc^ reporter mice with or without topical clobetasol treatment of the injection site (b). The AUC of the curves in (b) are shown in (c) (n = 4). Luciferase expression kinetics after intradermal electroporation of LUC-sa-mRNA in wild type BABL/c mice after silica- or cellulose-based purification with or without clobetasol treatment of the injection site (d). The AUC of the curves in (d) are shown in (e). Each symbol or bar represents the mean of four individual mice and the error bars represent SEM. The statistical analysis of the data shown in (b) and (d) can be found in Table S1.

### Cellulose-purified self-amplifying mRNA vaccines elicit a stronger humoral and cellular immune response

We finally investigated whether the novel cellulose-based purification method could improve the efficacy of our ZIKVac-sa-mRNA vaccine. It has been reported that inhaled and oral corticosteroids do not affect the efficacy of influenza vaccines [19, 20]. We were triggered by these contraintutives reports and therefore decided to also investigate the effect of clobetasol pre- and posttreatment on the efficacy of our ZIKVac-sa-mRNA vaccine. Forty-eight BALB/c mice were randomized in six groups and vaccinated by intradermal electroporation of either cellulose- or silica-purified ZIKVac-sa-mRNA with or without clobetasol. LUC-sa-mRNA was used as negative control (Figure 6a-b).

**Fig. 6.**
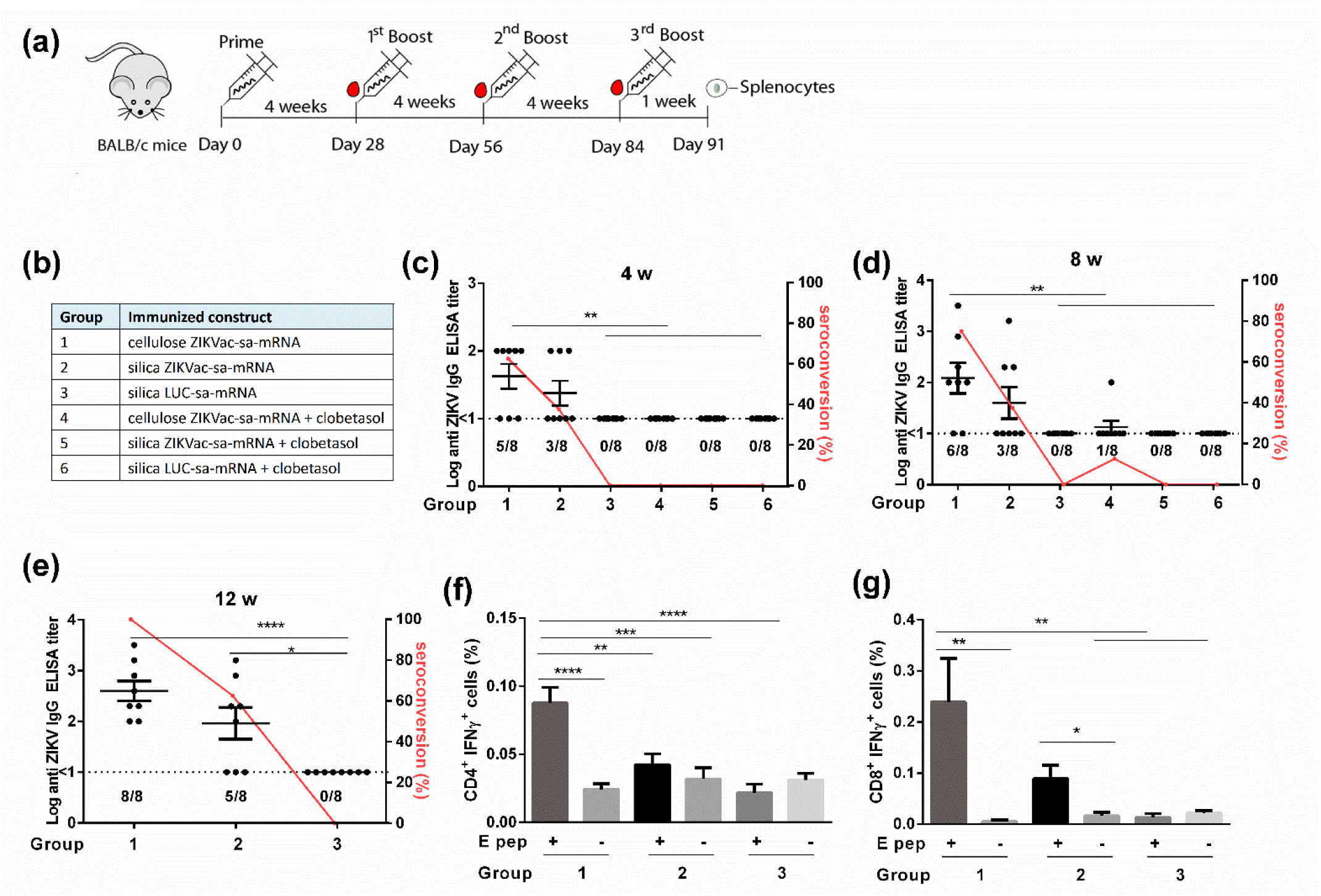
Vaccination efficacy of silica- and cellulose-purified ZIKVac-sa-mRNA in BALB/c mice topically treated with or without clobetasol. Experimental setup (a) and overview of the different groups (b). Mice were intradermally electroporated with 1 μg of celluose-or silica-purified ZIKVac-sa-mRNA vaccine or LUC-sa-mRNA control on day 0, day 28, day 56 and day 84 (groups 1-3, without clobetasol treatment) or on day 0 and day 28 (groups 4-6, with clobetasol treatment). Antibody titers in mice were determined with a ZIKV E-protein-specific IgG ELISA at 4 weeks (c), 8 weeks (d) or 12 weeks (e) after initial immunization (n = 8). The dashed lines indicate the limit of detection of the assay. The percentage of seroconversed mice is depicted with a red line (right Y-axis). Antigen-specific CD4^+^ (f) and CD8^+^ (g) T cell responses in mice from groups 1-3 were assessed one week after 3^rd^ boost by IFN-γ staining in T cells stimulated with a ZIKV E protein peptide pool (E pep +). Each bar represents the mean of eight individual mice and the error bars represent SEM.

The interval between the vaccinations was four weeks and the dose was 1 µg. Compared to the silica-purified vaccine, higher antibody titers and seroconversion rates were observed in mice receiving the cellulose-purified ZIKVac-sa-mRNA vaccine, both after the prime as well as after the boost vaccination (Figure 6c-d). As expected, a second vaccination further increased the antibody titers and seroconversion rates of both vaccines. In more detail, four weeks after the first boost the mean antibody titer and seroconversion rate of the mice vaccinated with the cellulose-purified ZIKVac-sa-mRNA vaccine was 122 and 75%, while these values were 40 and 37.5% in the mice that received the silica-purified ZIKVac-sa-mRNA vaccine (Fig. 6d). Topical application of clobetasol at the injection site completely abolished the efficacy of the ZIKVac-sa-mRNA vaccine (Figure 6c-d). As the humoral immune response improved after the boost vaccination, we decided to give a second boost to the mice that received the ZIKVac-sa-mRNA vaccine without co-administration of clobetasol (Group 1-3). Again, the antibody titer and seroconversion rates increased in both vaccinated groups and the mice that received the cellulose-purified sa-mRNA vaccine developed the highest antibody titers. Moreover, the seroconversion rate in this group increased to 100%, while the seroconversion rate in mice immunized with silica-purified ZIKVAc-sa-mRNA vaccine was only 62.5% (Figure 6e). In addition to this strong humoral response, increased ZIKV E protein-specific CD4^+^ and CD8^+^ T cell responses were seen after the final immunization with the cellulose-purified ZIKVac-sa-mRNA vaccine. These cellular responses were significantly higher than those obtained with LUC-sa-mRNA vaccinated mice (Figure 6f-g).

## Discussion

Synthetic self-amplifying mRNAs are known for their high *in vivo* translational efficiency [21, 22]. However, *in vivo* administration of sa-mRNAs may induce a strong type I IFN response [5, 11, 23]. While this can be considered advantageous when the sa-mRNA is used for vaccination purposes [11, 24], several studies have demonstrated that triggering type I IFN production can negatively impact the intended adaptive immune response of intramuscularly and intradermally administered mRNA vaccines [9, 11, 15]. Evidently, a strong type I IFN mediated inflammatory response should also be avoided when synthetic mRNAs are considered for protein-replacement therapy, gene editing or stem cell reprogramming [25, 26]. In line with previous reports [8, 11, 15], we found that intradermal electroporation of our formerly developed ZIKVac-sa-mRNA vaccine results in a very rapid upregulation of type I IFNs, with a maximal induction within 5 hours after sa-mRNA administration (Fig. 2). By-products originating from the in vitro transcription process like double strand (ds), uncapped or untailed RNAs species are contributing to this innate immune response [1]. There is also a concern that the intracellular amplification of sa-mRNAs, which occurs through dsRNA intermediates, may strongly trigger intracellar sensors such as RIG-I and MDA5 [27]. However, our data do not support this idea, as the peak in IFN-β production occurs directly after the delivery of the sa-mRNA and thus not at the moment of sa-mRNA replication. Moreover, we recently reported a similar IFN-β induction for replication-deficient and replication-competent sa-mRNAs in mice [15].

In an attempt to block the immediate type I IFN response elicited by our ZIKVac-sa-mRNA vaccine, we screened several commercial TLR and NF-κB/NLRP3 inhibitors (Fig. 1). Endosomal or cell surface associated TLRs are one type of PRRs that recognize dsRNAs, uncapped or untailled ssRNAs by-products present in synthetic sa-mRNA [28]. Co-injection of TLR7 (ODN20958) or TRL7/8/9 (ODN2088) antagonists with the ZIKVac-sa-mRNA vaccine seems to shortly block the recognition of ssRNA species (Fig. 2). This indicates that TLR7 and/or 8, which recognize single nucleosides and short ssRNAs (oligoribonucleotides), are involved in sensing our sa-mRNA vaccine [7, 29]. We hypothesize that also degradation products occurring from sa-mRNAs that did not enter the cells after *in vivo* electroporation are recognized by cell surface associated TLR7/8. Co-administration of our sa-mRNA vaccine with BAY11, which blocks nuclear translocation of NF-κB and inhibits the NLRP3 inflammasome [30, 31], also significantly decreased the initial innate immune response. The short-lived downregulation of the type I IFN response by co-injected ODN2088, ODNA20958 or BAY11 is probably due to a rapid dilution of the inhibitors from the injection site. Therefore, we tested pre- and post-treatment of the injection site with these inhibitors. However, this only slightly increased and prolonged the reduction of the innate immune response (Fig. 2). Probably, pre- and posttreatment with these inhibitors should be performed closer to the moment of injection, e.g. 15 min (instead of 5 h) before and 1 h (instead of 12 h) after injection of the sa-mRNA vaccine. An interesting future approach would be the co-encapsulation of these innate immune inhibitors with sa-mRNA into e.g. lipid nanoparticles. Interestingly, no diminution of the type I IFN response was observed when 12.5 µg or even 25 µg of a TLR3 inhibitor was co-administrated (Fig. S1). This is remarkable, since a dot blot assay clearly indicated that dsRNA species are present in the silica-purified sa-mRNA (Fig. 5a). Possibly the massive amounts of dsRNA in our silica-purified sa-mRNA completely competed out the binding of the TLR3 antagonist to the TLR3 recepetors.

Besides the aforementioned specific inhibitors of the innate immune response, we also tested clobetasol propionate, which is a potent corticosteroid with a broad mode of action [32, 33]. Topical application of clobetasol at the injection site efficiently inhibited the type I IFN response elicited by our ZIKVac-sa-mRNA vaccine. However, it is essential that the injection spot is pretreated with clobetasol prior to sa-mRNA administration (Fig. 2 and 3). Moreover, additive effects were observed when topical clobetasol was combined with ODN2088 and/or BAY11 (Fig. 3). However, only clobetasol was able to significantly improve the *in vivo* translation efficacy of sa-mRNA (Fig. 4 and Fig. S2). A combination of pre-, co- and post-treatment of the injection site with clobetasol prolonged the expression of the luciferase encoding sa-mRNA with one week and significantly increased the overall expression (Fig. 4 and Fig. S2c). Surprisingly, ODN2088 did not improve the expression and BAY11 even decreased the *in vivo* expression of the sa-mRNA (Fig. 4 and Fig. S2a-b). This observation corresponds with the findings of Liu et al. (2017), who screened 15 different inhibitors and found that reduced IFN production was not associated with enhanced mRNA translation in cultured human foreskin fibroblast cells [34]. Similar to our results, 7 of the tested inhibitors even reduced the mRNA translation efficacy [34]. In contrast, Awe et al. (2013) reported enhanced *in vitro* translation of the transcription factor OCT4 from a synthetic mRNA upon BAY11 supplementation [35]. However, the enhanced OCT4 translation was not achieved when the type I IFN decoy receptor B18R was supplemented, indicating that the observed increase in translation was independent of type I interferons [35].

As mentioned earlier, synthetic mRNAs produced by *in vitro* transcription contain by-products such as dsRNAs and small abortive ssRNA species which are known to strongly stimulate innate immune responses in mammalian cells [1, 18]. Cellulose-purification is reported to efficiently remove small by-products like free ribonucleotides and dsRNA species larger than 30 bp [1, 18, 36]. The method has been successfully applied on non-amplifying in vitro transcribed mRNA [18]. Here we investigated whether cellulose-based purification [18] could also reduce the type I IFN response and increase the *in vivo* translation efficacy of sa-mRNAs by removing dsRNA by-products. Immunoblotting confirmed that cellulose-mediated purification of sa-mRNA efficiently removed dsRNA species and, compared to the standard silica-based purification, the cellulose-purified sa-mRNA elicited a much lower type I IFN response, which could be further reduced by topical clobetasol (Fig. 5). However, this beneficial effect of cellulose purification was not completely reflected in the translation efficacy, since significant increases in translation efficacy were only observed when the mice were treated with clobetasol (Fig. 5). The UTRs in our sa-mRNA are based on the RNA genome of VEEV, which posess structural elements in its 5-UTR that can (partly) evade translational inhibition induced by a type I IFN response [37]. Therefore, this may explain why strategies that reduce the type I IFN do not drastiscally improve the translation efficacy of our synthetic sa-mRNA. Alternatively, it is also possible that the drop in type I IFNs induced by TLR7/8 antagonists or cellulose-purification is not enough to improve the translation efficiency of the sa-mRNA. To further decrease the type I IFN response, inhibitors for other PRRs like RIG-I like receptor can be used and the cellulose-based purification can be further improved. Indeed, the dot blot (Fig. 5a) show some remaining dsRNA species. Baiersdörfer et al. reported that these are remaining dsRNAs are mainly short (<30 bp) dsRNAs [18]. Short uncapped dsRNAs are especially recognized by RIG-I [38]. Therefore, phosphatase treatment of the sa-mRNA prior to injection can also circumvent RIG-I-mediated detection of di/tri-phosphate 5’ ends [39].

In a final experiment, we demonstrated that the cellulose-purified ZIKVac-sa-mRNA vaccine induced higher antigen specific humoral and cellular immune responses than the silica-purified ZIKVac-sa-mRNA vaccine. This confirms that the by-products after in vitro transcription exert negative effects on the efficacy of our sa-mRNA-based vaccine. These results also support our previous findings that silica-purified ZIKVac-sa-mRNA elicits stronger humoral and cellular immune responses in IFNAR1^-/-^ mice, which show defective type I IFN signaling [15]. Clobatesol treatment of the vaccination site prevented the induction of a humoral immune response, despite of the beneficial effects of clobetasol on the IFN response and the translation of the sa-mRNA (Fig. 6). This is an important finding as it was reported that inhaled and oral corticosteroids do not affect the efficacy of influenza vaccines [19, 20]. Moreover, these data indicate that corticosteroids can be used to prevent that antibodies are raised against mRNA encoded therapeutic proteins like e.g. clotting factors or erythropoietin.

In summary, among a handful of commercial TLR and NF-κB/NLRP3 inhibitors, topical application of clobetasol caused the strongest reduction of the innate immune response elicited after intradermal electroporation of our ZIKVac-sa-mRNA vaccine. Combining clobetasol with a TLR7 antagonist and/or a NRLP-3/NF-κB inhibitor further reduced the innate immune response. Clobetasol also increased the translation of intradermally electroporated sa-mRNA. In a vaccination context, however, co-administration of clobetasol with our ZIKVac-sa-mRNA vaccine completely blocked the cellular and humoral immune response. In contrast, purification of the ZIKVac-sa-mRNA vaccine with a novel cellulose-based method tripled the antibody titers, doubled the cellular immune response an increased the seroconversion rate from 62.5% to 100%. This improvement was associated with a drastic reduction of dsRNA by-products, which significantly decreased the type I intereferon response elicited by the sa-mRNA vaccine and slightly improved the expression of the sa-mRNA. It is important to note that the data in this study were achieved by intradermal electroportion of the sa-mRNA vaccine and thus without the use of a carrier. In a future project we aim to determine if this novel purification method also improves the efficacy of sa-mRNAs that are delivered by lipid nanoparticles.

## Materials and Method

### Mice

Female BALB/c mice, aged 6-8 weeks, were purchased from Janvier (France). Heterozygous albino (tyr^c2J^) C57BL/6 IFN-β reporter (IFN-β^+/Δβ-luc^) mice used in this study were from the Institute for Laboratory Animal Science, Hannover Medical School (Germany) and the breed was further maintained in house. All mice were housed in individually ventilated cages and had free access to feed and water. Mice experiments were approved by the Ethics Committee of the Faculty of Veterinary Medicine, Ghent University (No. EC2019/62). During intradermal injections and bioluminescence imaging, mice were under isoflurane anesthesia (5% for induction and 2% for maintenance).

### Inhibitors

TLR3 and 7 inhibitors ODN2088 and ODN20958 (Miltenyi Biotech, Belgium) were used in this study. The phosphorothioate modified ODN2088 (5’-TCCTGGCGGGGAAGT-3’) is a TLR7/8/9 antagonist, while the phosphorothioate modified oligonucleotide ODN20958 is a TLR-7 inhibitor (5’-TCCTAACAAAAAAAT-3’). The TLR3/dsRNA complex inhibitor (C_18_H_13_ClFNO_3_S) was bought from Merck Millipore (Belgium). The NF-κB and NOD-like receptor pyrin 3 (NLRP3) inhibitor BAY11-7082 was from Invivogen (Belgium). The TLR3/dsRNA complex inhibitor and BAY11-7082 were dissolved in DMSO. Clobetasol propionate ointment (0.05%, Dermovate™ Cream) was from GlaxoSmithKline (GSK).

### mRNA and silica purification

Self-amplifying mRNA (sa-mRNA) was synthetized via *in vitro* transcription (IVT) as described [15]. Briefly, ZIKVac-sa-mRNA was constructed by inserting the sequence of the Zika virus prM-E fusion protein of the Brazilian Rio-S1 ZIKV strain (GenBank No. KU926310.1) containing a signal peptide of Japanese encephalitis virus (JEV) at the 5’ terminal end into the pTK155 plasmid using Gateway Cloning (Invitrogen). The sequence of firefly luciferase (LUC) was cloned into pTK155 to produce LUC-sa-mRNA. The plasmids of VEEV-based ZIKVac-sa-mRNA and LUC-sa-mRNA were transformed into competent *E. coli* bacteria (Invitrogen, Massachusetts, USA) and after 24 h purified with the Plasmid Plus Midi kit (Qiagen, Germany). Subsequently, linearized plasmids were obtained using I-SceI endonuclease (NEB, Massachusetts, USA) and the sa-mRNAs were synthetized by IVT with a MEGAscript T7 Transcription kit (Life Technologies, Massachusetts, USA). Next the sa-mRNA was purified with the RNeasy^®^ Mini kit (Qiagen, Germany) and post-transcriptionally capped using a ScriptCap m7G Capping System and the 2’-O-Methyltransferase kit (Cellscript, Wisconsin, USA) to obtain cap-1. After capping, the sa-mRNA was purified again with the RNeasy^®^ Mini kit (Qiagen, Germany). As a 40-nucleotide-long poly(A) was encoded in the linearized plasmid template, Poly(A) tailing was not required. The quantity and quality of the sa-mRNAs were determined with a Nanodrop spectrophotometer (Thermo Fisher Scientific, Massachusetts, USA) and sa-mRNAs were stored at −80 °C.

### Cellulose-based purification of sa-mRNAs

After IVT the mRNAs (LUC-sa-mRNA and ZIKVac-sa-mRNA) were purified by LiCl precipitation and subsequently enzymatically capped as described above. Next, the capped sa-mRNAs were again precipitated with LiCl and resuspended in HEPES-ethanol buffer (10 mM HEPES, pH7.2, 0.1 mM EDTA, 125 mM NaCl and 16% ethanol). Subsequently, an additional cellulose-based purification was performed to remove dsRNA by-products as previously described [18]. Briefly, cellulose fibers (Sigma-Aldrich, Belgium) were suspended in HEPES-ethanol buffer at a concentration of 0.2 g/mL. After 10 min of vigorous mixing, 630 µl of the cellulose suspension was transferred to a microcentrifuge spin column (NucleoSpin Filters, Macherey-Nagel, Düren, Germany) and centrifuged for 1 min at 14,000 g. The flow through was discarded and 450 µl HEPES-ethanol buffer was added to the cellulose fibers followed by vigorously shaking for 5 min. Subsequently, the spin column was centrifuged for 1 min at 14,000 g and the flow through was discarded. Hundred to 500 μg of sa-mRNA in 450 μl HEPES-ethanol buffer was added to the cellulose in the spin column followed by vigorously shaking for 30 min to allow association of the dsRNA by-products to the cellulose. Separation of the cellulose associated dsRNA from the sa-mRNA occurred by centrifugation at 14,000 g for 1 min. The collected flow through containing the sa-mRNA was precipitated by adding 0.1 volume of 3 M NaOAc pH 5.5 (50 μl) and 1 volume of isopropanol and incubating this mixture for 30 min at -20 °C. Next, the mRNA was pelleted by centrifugation at 4 °C for 15 min at 14,000 g and the supernatant was discarded. The pellet was washed with 500 μl 70% pre-cooled ethanol and centrifuged at 4 °C for 5 min at top speed. The supernatant was removed and the cellulose-purified mRNA was finally dissolved in nuclease-free water.

### Dot blot analysis of dsRNA by-products

Silica- or cellulose-purified sa-mRNAs (LUC-sa-mRNA and ZIKVac-sa-mRNA) were diluted in nuclease-free water to final concentrations of 40 and 200 ng/μl. Subsequently 5 μl aliquots (200 or 1000 ng sa-mRNAs per dot) were loaded to a positively charged nylon membrane (Whatman Nytran SuPerCharge, Sigma-Aldrich) that was tapped on a sheet of Whatman GB005 blotting paper. After drying, the membrane was blocked in 5% (w/v) non-fat dried milk in PBS-T buffer (0.1% (v/v) Tween-20 in PBS) for 1 h at room temperature. After three washes with PBS-T buffer, the membrane was incubated overnight at 4°C on a rolling mixer with mouse J2 anti-dsRNA murine antibody (Scicons, Budapest, Hungary) diluted 1:5,000 in PBS-T buffer containing 1% (w/v) non-fat dried milk. Next, the membranes were washed three times with PBS-T buffer before and incubated for 1h at room temperature with horseradish peroxidase (HRP)-conjugated donkey anti-mouse IgG (H+L, Jackson ImmunoResearch Laboratories, Cambridgeshire, UK) diluted 1:10,000 in PBS-T buffer containing 1% (w/v) non-fat dried milk. After washing the membranes three times with PBS-T buffer, the detection of the target dsRNAs on the membrane was performed using SuperSignal West Femto Maximum Sensitivity Substrate (Thermo Scientific) and the ChemiDoc MP Imaging System (Bio-Rad, USA).

### In vivo interferon response

Interferon beta reporter (IFN-β^+/Δβ-luc^) mice were used to investigate the effect of several innate immune inhibitors on the interferon response elicted by our ZIKVac-sa-mRNA vaccine. IFN-β^+/Δβ-luc^ mice were shaved at their flanks and intradermally injected at both flanks with 0.5 µg sa-mRNA vaccine in 25 µl PBS (without Ca^2+^ and Mg^2+^) using 29 G insulin needles (VWR, Netherlands). Electroporation, when used, was performed immediately after each sa-mRNA injection with a 2-needle array electrode containing 4 needles per row of 4 mm (AgilePulse, BTX Harvard Apparatus, Massachusetts, USA). The procedure of electroporation involved two short high-voltage pulses of 450 V with a duration of 0.05 ms and an interval of 300 ms followed by eight long low-voltage pulses of 100 V with a duration of 10 ms and an interval of 300 ms [4, 15]. Innate immune inhibitors (Fig. 1) were co-injected with the sa-mRNA vaccine. In certain experiments the injection site was 5 h before sa-mRNA administration also pretreated and posttreated twice daily by intradermal injection of the innate inhibitors. The corticosteroid clobetasol was not injected but topically applied as an ointment at the injection site (25 µg clobetasol per cm^2^) and in certain experiments pre- and posttreatments was also evaluated. The type I IFN response was monitored by measuring daily during 7 days the bioluminescent signal at the injection sites. To that end, mice were subcutaneously injected with 200 µl D-luciferin (15 mg/ml, Gold Biotechnology, USA). Twelve minutes later the mice were anesthesized using isoflurane and the in vivo bioluminescence signal was recorded using an IVIS Lumina II (PerkinElmer, USA).

### In vivo translation kinetics

The effect of selected innate inhibitors and the cellulose-based purification method on the *in vivo* translation of the sa-mRNA was investigated by intradermal electroporation (see above for protocol) of 1 µg silica- or cellulose-purified LUC-sa-mRNA in BALB/c mice in the presence or absence of the indicated innate immune inhibitors. The luciferase expression was monitored during 28 or 35 days by *in vivo* optical imaging as described above.

### Vaccination experiment

Preshaved female BALB/c mice (6 weeks) were anesthetized by inhalation of isoflurane and immunized by intradermal electroporation of 0.5 µg ZIKVac-sa-mRNA vaccine or the LUC-sa-mRNA in both flanks using the vaccination schedule depicted in Fig. 6a. Silica- and cellulose-purified ZIKVac-sa-mRNA vaccines with and without treatment of the injection site with clobetasol were investigated. Topical treatment of clobetasol involved a pretreatment of the injection site 12 h prior to vaccination and a posttreatment twice daily until 3 days after immunization. Electroporation was performed immediately after each ZIKVac-sa-mRNA injection using the protocol described above.

### Zika virus specific antibody titers

The mouse ZIKV ELISA kit (Alpha Diagnostic International, TX, USA) was used to determine ZIKV E protein-specific antibody titers. In more details, 96-well plates that were pre-coated with ZIKV E protein were equilibrated for 5 min at room temperature with 300 μl of the provided wash buffer. Subsequently, two-fold serial dilutions of the serum samples were made (starting from a 50-fold dilution) and 100 µl of these dilutions was added per well along with the calibration standards. After 1 h of incubation at room temperature, the plates were washed four times with wash solution. Next, 100 μl of anti-mouse IgG HRP-conjugate working solution was added to the wells and incubated at room temperature. After 30 min, the wells were washed five times and subsequently incubated with 100 μl of 3,3’,5,5’-Tetramethylbenzidine (TMB) substrate at room temperature. The enzymatic conversion of TMB was stopped after 15 min by adding 100 μl of stop solution and the absorbance was measured at 450 nm in a Biochrom EZ 400 microplate reader (Biochrom, England). The antibody endpoint titers were defined as the highest reciprocal dilution with an absorbance that was at least two times the background (obtained with serum of unvaccinated mice).

### Zika Virus specific cellular immune response

Intracellular cytokine staining was performed to determine Zika virus specific CD4^+^ and CD8^+^ T cells responses with flow cytometry. In more detail, splenocytes were isolated one week after the last boost and stimulated in 96-well plates (1×10^6^ cells/well) with 2 μg/ml of overlapping 15-amino-acid peptides covering the ZIKV E protein (JPT, Berlin, Germany) in 1640 RPMI medium. After 1 h of stimulation at 37 °C 0.3 µl of eBioscience(tm) protein transport inhibitor cocktail (Brefeldin A 5.3 mM + Monensin 1 mM, eBioscience) was added to 150 µl of stimulated splenocytes and the samples were further incubated for 5 h at 37 °C. Splenocytes were then harvested, washed with cold PBS, treated with mouse BD Fc Block(tm) (BD Biosciences), and stained with anti-CD3-APC/CD4-PerCP/CD8-Alexa Fluor 488 antibodies (clones 145-2C11, RM4-5 and 53-6.7, Biolegend) for 30 min at 4 °C according to the manufacturer’s instructions. Subsequently, the cells were fixed and permeabilized with the Fixation/Permeabilization buffer (eBioscience) for 30 min at 4 °C before intracellular staining with anti-IFN-γ-PE antibody (clone XMG1.2, Biolegend) for 30 min at room temperature. All the samples were finally washed and stored at 4 °C until analysis using a Cytoflex flow cytometer (Beckman Coulter). Single and live cells were gated and 300,000 events were collected for each sample. Samples treated with Cell Stimulation Cocktail (eBioscience) served as positive controls and unstimulated samples as negative controls.

### Statistical analyses

Statistical analyses were performed with GraphPad Prism software (version 7.0, GraphPad Software Inc., CA, USA). The longitudinal experiments of different animal groups were analyzed using repeated-measures two-way ANOVA, corrected for multiple comparisons (Bonferroni method). Differences between two groups were compared with student’s t test (non-parametric Mann-Whitney U test). Data in the study are represented as means ± SEM, unless otherwise mentioned. A p-value of below 0.05 is considered statistically significant difference (*p<0.05, **p<0.01, ***p<0.001,

****p<0.0001).

## Acknowledgements

This work was supported by the concerted research action (GOA) found of Ghent University (project code BOF15/GOA/013) and the research foundation-Flanders (FWO, project code G087516N). Z.Z. acknowledges the funding from China Scholarship Council (CSC) (201607650018).

## Author Contributions

Z.Z. and N.N.S. conceived experiments, participated in experimental studies, interpreted the results and wrote the manuscript. S.M.C and L.O. assisted to perform cellulouse-based purification of sa-mRNA. F.C. and N.N.S. critically revised the manuscript. S.M.C, H.W, H.H, J.D.T and J.P.P.C helped with the *in vivo* experiments. S.L. provided IFN-β^+/Δβ-luc^ reporter mice.

All authors have given approval to the final version of the manuscript.

---

**Fig. S1.**
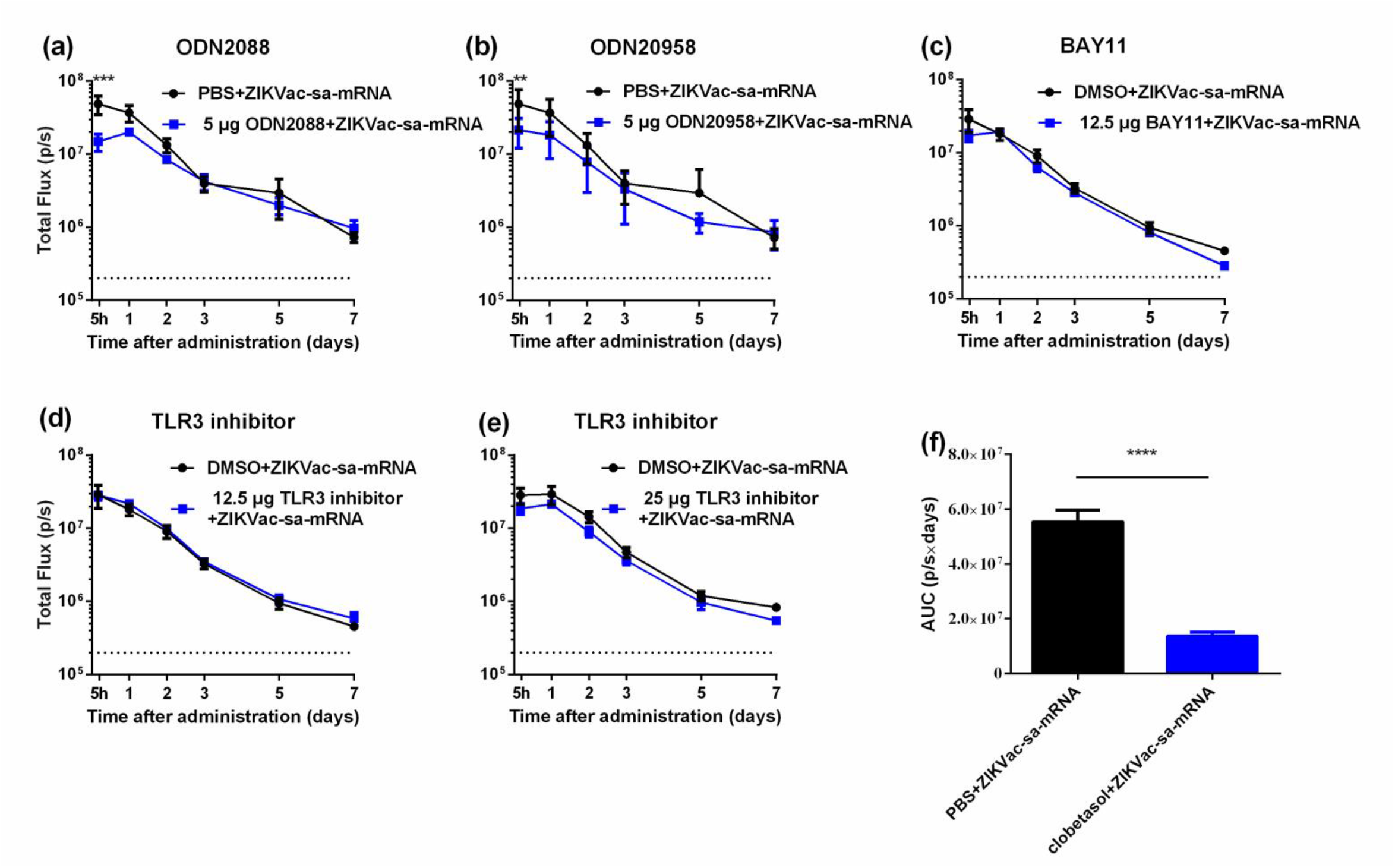
Effect of the TLR3 inhibitor and lower doses of ODN2088, ODN20958 and BAY11 on the type I IFN response after intradermal electroporation of ZIKVac-sa-mRNA in IFN-β^+/Δβ-luc^ reporter mice. The effects of 5 μg ODN2088 (a), 5 μg ODN20958 (b), 12.5 μg BAY11 (c), 12.5 μg and 25 μg TLR3 inhibitor (d-e) on the type I IFN response of intradermally electroporated ZIKVac-sa-mRNA (1 µg). These inhibitors were mixed with the ZIKVac-sa-mRNA upon administration. (n = 4, data are shown as means ± SEM). In addition, the area under the curves (AUC) of Fig. 2h is presented in (f).

**Fig. S2.**
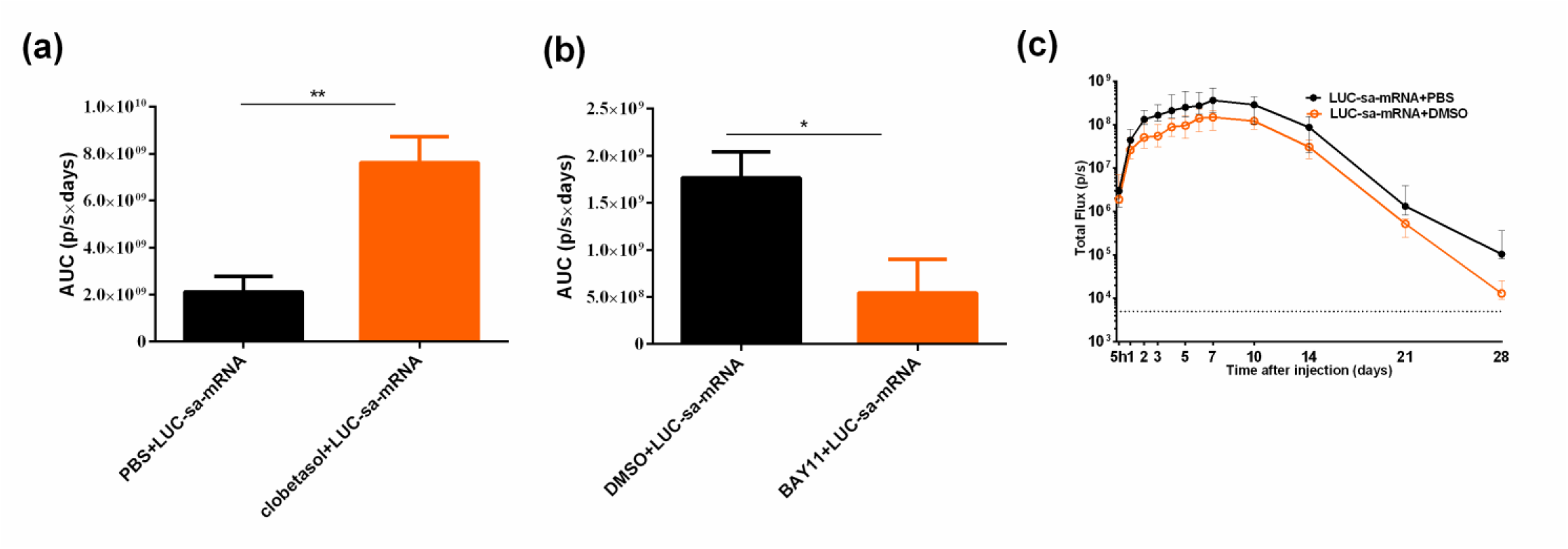
The influence of innate immune inhibitors on the translation of LUC-sa-mRNA in BALB/c mice. The area under the curves (AUCs) of Fig. 4b and 4d are presented in (a) and (b), respectively. Each bar represents the mean of four individual mice and the error bars represent SEM. The luciferase expression was determined in BABL/c mice that were intradermally electroporated with 1 µg of LUC-sa-mRNA with DMSO or PBS, each symbol represents the median of four individual mice and the error bars represent interquartile range (c).

