## Supplemental Table 1 for "Corticosteroids and cellulose purification improve respectively the in vivo translation and vaccination efficacy of self-amplifying mRNAs"

| **Table S1. Significant difference for Figure 5b and Figure 5d.** | | | |
| --- | --- | --- | --- |
| *Significant difference for Figure 5b* | | | |
| **5 h** |  | **Sinificance** | **P value** |
| silic ZIKVac-sa-mRNA+clobetasol | Cellulose ZIKVac-sa-mRNA+clobetasol | ns | 0.2241 |
| silic ZIKVac-sa-mRNA+clobetasol | Silic ZIKVac-sa-mRNA | **** | < 0.0001 |
| silic ZIKVac-sa-mRNA+clobetasol | Cellulose ZIKVac-sa-mRNA | ns | > 0.9999 |
| Cellulose ZIKVac-sa-mRNA+clobetasol | Silic ZIKVac-sa-mRNA | **** | < 0.0001 |
| Cellulose ZIKVac-sa-mRNA+clobetasol | Cellulose ZIKVac-sa-mRNA | ns | 0.7716 |
| Silic ZIKVac-sa-mRNA | Cellulose ZIKVac-sa-mRNA | **** | < 0.0001 |
| **1 day** |  | **Sinificance** | **P value** |
| silic ZIKVac-sa-mRNA+clobetasol | Cellulose ZIKVac-sa-mRNA+clobetasol | ns | > 0.9999 |
| silic ZIKVac-sa-mRNA+clobetasol | Silic ZIKVac-sa-mRNA | **** | < 0.0001 |
| silic ZIKVac-sa-mRNA+clobetasol | Cellulose ZIKVac-sa-mRNA | ns | > 0.9999 |
| Cellulose ZIKVac-sa-mRNA+clobetasol | Silic ZIKVac-sa-mRNA | **** | < 0.0001 |
| Cellulose ZIKVac-sa-mRNA+clobetasol | Cellulose ZIKVac-sa-mRNA | ns | > 0.9999 |
| Silic ZIKVac-sa-mRNA | Cellulose ZIKVac-sa-mRNA | **** | < 0.0001 |

| *Significant difference for Figure 5d* |  |  |  |
| --- | --- | --- | --- |
| **2 days** |  | **Sinificance** | **P value** |
| Silic LUC-sa-mRNA + clobetasol | Cellulose LUC-sa-mRNA + clobetasol | ns | > 0.9999 |
| Silic LUC-sa-mRNA + clobetasol | Silic LUC-sa-mRNA | **** | < 0.0001 |
| Silic LUC-sa-mRNA + clobetasol | Cellulose LUC-sa-mRNA | **** | < 0.0001 |
| Cellulose LUC-sa-mRNA + clobetasol | Silic LUC-sa-mRNA | **** | < 0.0001 |
| Cellulose LUC-sa-mRNA + clobetasol | Cellulose LUC-sa-mRNA | *** | 0.0003 |
| Silic LUC-sa-mRNA | Cellulose LUC-sa-mRNA | ns | > 0.9999 |
| **3 days** |  | **Sinificance** | **P value** |
| Silic LUC-sa-mRNA + clobetasol | Cellulose LUC-sa-mRNA + clobetasol | ** | 0.0093 |
| Silic LUC-sa-mRNA + clobetasol | Silic LUC-sa-mRNA | *** | 0.0001 |
| Silic LUC-sa-mRNA + clobetasol | Cellulose LUC-sa-mRNA | ns | 0.0958 |
| Cellulose LUC-sa-mRNA + clobetasol | Silic LUC-sa-mRNA | **** | < 0.0001 |
| Cellulose LUC-sa-mRNA + clobetasol | Cellulose LUC-sa-mRNA | **** | < 0.0001 |
| Silic LUC-sa-mRNA | Cellulose LUC-sa-mRNA | ns | 0.9991 |
| **4 days** |  | **Sinificance** | **P value** |
| Silic LUC-sa-mRNA + clobetasol | Cellulose LUC-sa-mRNA + clobetasol | ** | 0.0081 |
| Silic LUC-sa-mRNA + clobetasol | Silic LUC-sa-mRNA | *** | 0.0001 |
| Silic LUC-sa-mRNA + clobetasol | Cellulose LUC-sa-mRNA | * | 0.0235 |
| Cellulose LUC-sa-mRNA + clobetasol | Silic LUC-sa-mRNA | **** | < 0.0001 |
| Cellulose LUC-sa-mRNA + clobetasol | Cellulose LUC-sa-mRNA | **** | < 0.0001 |
| Silic LUC-sa-mRNA | Cellulose LUC-sa-mRNA | ns | > 0.9999 |
| **5 days** |  | **Sinificance** | **P value** |
| Silic LUC-sa-mRNA + clobetasol | Cellulose LUC-sa-mRNA + clobetasol | ** | 0.0015 |
| Silic LUC-sa-mRNA + clobetasol | Silic LUC-sa-mRNA | ns | 0.3438 |
| Silic LUC-sa-mRNA + clobetasol | Cellulose LUC-sa-mRNA | ns | > 0.9999 |
| Cellulose LUC-sa-mRNA + clobetasol | Silic LUC-sa-mRNA | **** | < 0.0001 |
| Cellulose LUC-sa-mRNA + clobetasol | Cellulose LUC-sa-mRNA | **** | < 0.0001 |
| Silic LUC-sa-mRNA | Cellulose LUC-sa-mRNA | ns | > 0.9999 |
| **6 days** |  | **Sinificance** | **P value** |
| Silic LUC-sa-mRNA + clobetasol | Cellulose LUC-sa-mRNA + clobetasol | ns | 0.7251 |
| Silic LUC-sa-mRNA + clobetasol | Silic LUC-sa-mRNA | ns | 0.6574 |
| Silic LUC-sa-mRNA + clobetasol | Cellulose LUC-sa-mRNA | ns | > 0.9999 |
| Cellulose LUC-sa-mRNA + clobetasol | Silic LUC-sa-mRNA | *** | 0.0001 |
| Cellulose LUC-sa-mRNA + clobetasol | Cellulose LUC-sa-mRNA | * | 0.0205 |
| Silic LUC-sa-mRNA | Cellulose LUC-sa-mRNA | ns | > 0.9999 |
| **7 days** |  | **Sinificance** | **P value** |
| Silic LUC-sa-mRNA + clobetasol | Cellulose LUC-sa-mRNA + clobetasol | ** | 0.0087 |
| Silic LUC-sa-mRNA + clobetasol | Silic LUC-sa-mRNA | ns | > 0.9999 |
| Silic LUC-sa-mRNA + clobetasol | Cellulose LUC-sa-mRNA | ns | > 0.9999 |
| Cellulose LUC-sa-mRNA + clobetasol | Silic LUC-sa-mRNA | **** | < 0.0001 |
| Cellulose LUC-sa-mRNA + clobetasol | Cellulose LUC-sa-mRNA | **** | < 0.0001 |
| Silic LUC-sa-mRNA | Cellulose LUC-sa-mRNA | ns | > 0.9999 |
| **14 days** |  | **Sinificance** | **P value** |
| Silic LUC-sa-mRNA + clobetasol | Cellulose LUC-sa-mRNA + clobetasol | ns | 0.9238 |
| Silic LUC-sa-mRNA + clobetasol | Silic LUC-sa-mRNA | ns | 0.6629 |
| Silic LUC-sa-mRNA + clobetasol | Cellulose LUC-sa-mRNA | ns | 0.3951 |
| Cellulose LUC-sa-mRNA + clobetasol | Silic LUC-sa-mRNA | *** | 0.0005 |
| Cellulose LUC-sa-mRNA + clobetasol | Cellulose LUC-sa-mRNA | *** | 0.0001 |
| Silic LUC-sa-mRNA | Cellulose LUC-sa-mRNA | ns | > 0.9999 |
